# Cell Population Growth Kinetics in the Presence of Stochastic Heterogeneity of Cell Phenotype

**DOI:** 10.1101/2023.02.08.527773

**Authors:** Yue Wang, Joseph X. Zhou, Edoardo Pedrini, Irit Rubin, May Khalil, Hong Qian, Sui Huang

**Author notes:** For correspondence (SH).

## Abstract

Recent studies at individual cell resolution have revealed phenotypic heterogeneity in nominally clonal tumor cell populations. The heterogeneity affects cell growth behaviors, which can result in departure from the idealized exponential growth. Here we measured the stochastic time courses of growth of an ensemble of populations of HL60 leukemia cells in cultures, starting with distinct initial cell numbers to capture the departure from the exponential growth model in the initial growth phase. Despite being derived from the same cell clone, we observed significant variations in the early growth patterns of individual cultures with statistically significant differences in growth kinetics and the presence of subpopulations with different growth rates that endured for many generations. Based on the hypothesis of existence of multiple inter-converting subpopulations, we developed a branching process model that captures the experimental observations.

## Introduction

Cancer has long been considered a genetic disease caused by oncogenic mutations in somatic cells that confer a proliferation advantage. According to the clonal evolution theory, accumulation of random genetic mutations produces cell clones with cancerous cell phenotype. Specifically, cells with the novel genotype(s) may display increased proliferative fitness and gradually outgrow the normal cells, break down tissue homeostasis and gain other cancer hallmarks (***Hanahan and Weinberg, 2011***). In this view, a genetically distinct clone of cells dominates the cancer cell population and is presumed to be uniform in terms of the phenotype of individual cells within an isogenic clone. In this traditional paradigm, non-genetic phenotypic variation within one clone is not taken into account.

With the advent of systematic single-cell resolution analysis, however, non-genetic cell heterogeneity within clonal (cancer) cell populations is found to be universal (***Pisco and Huang, 2015***). This feature led to the consideration of the possibility of biologically (qualitatively) distinct (meta)stable cell subpopulations due to gene expression noise, representing intra-clonal variability of features beyond the rapid random micro-fluctuations. Hence, transitions between the subpopulations, as well as heterotypic interactions among them may influence cell growth, migration, drug resistance, etc. (***Tabassum and Polyak, 2015; Gunnarsson et al., 2020; Durrett et al., 2011***). Thus, an emerging view is that cancer is more akin to an evolving ecosystem (***Gatenby et al., 2014***) in which cells form distinct subpopulations with persistent characteristic features that determine their mode of interaction, directly or indirectly via competition for resources (***Egeblad et al., 2010; Sonnenschein and Soto, 2000***). However, once non-genetic dynamics is considered, cell *“*ecology*”* differs fundamentally from the classic ecological system in macroscopic biology: the subpopulations can reversibly switch between each other whereas species in an ecological population do not convert between each other (***Clark, 1991***). This affords cancer cell populations a remarkable heterogeneity, plasticity and evolvability, which may play important roles in their growth and in the development of resistance to treatment (***Meacham and Morrison, 2013***).

Many new questions arise following the hypothesis that phenotypic heterogeneity and transitions between phenotypes within one genetic clone are important factors in cancer. Can tumors arise, as theoretical considerations indicate, because of a state conversion (within one clone) to a phenotype capable of faster, more autonomous growth as opposed to acquisition of a new genetic mutation that confers such a selectable phenotype (***Zhou et al., 2014a; Angelini et al., 2022; Howard et al., 2018; Sahoo et al., 2021; Pisco and Huang, 2015; Zhou et al., 2014b; Kochanowski et al., 2021***)? Is the macroscopic, apparently sudden outgrowth of a tumor driven by a new fastestgrowing clone (or subpopulation) taking off exponentially, or due to the cell population reaching a critical mass that permits positive feedback between its subpopulations that stimulates outgrowth, akin to a collectively autocatalytic set (***Hordijk et al., 2018***)? Should therapy target the fastest growing subpopulations, or target the interactions and interconversions of cancer cells?

At the core of these deliberations is the fundamental question on the mode of tumor cell population growth that now must consider the influence of inherent phenotypic heterogeneity of cells and the non-genetic (hence potentially reversible) inter-conversion of cells between the phenotypes that manifest various growth behaviors and the interplay between these two modalities. Traditionally tumor growth has been described as following an exponential growth law, motivated by the notion of uniform cell division rate for each cell, i.e. a first order growth kinetics (***Mackillop, 1990***). But departure from the exponential model has long been noted. To better fit experimental data, two major modifications have been developed, namely the Gompertz model and the West law model (***Yorke et al., 1993***). While no one specific model can adequately describe any one tumor, each model highlights certain aspects of macroscopic tumor kinetics, mainly the maximum size and the change in growth rate at different stages. These models however are not specifically motivated by cellular heterogeneity. Assuming non-genetic heterogeneity with transitions between the cell states, the population behavior is influenced by many intrinsic and extrinsic factors that are both variable and unpredictable at the single-cell level. Thus, tumor growth cannot be adequately captured by a deterministic model, but a stochastic cell and population level kinetic model is more realistic.

Using stochastic processes in modeling cell growth via clonal expansion has a long history (***Zheng, 1999***). An early work is the ***Luria and Delbrück*** (***1943***) model, which assumes cells grow deterministically, with wildtype cells mutating and becoming (due to rare and quasi-irreversible mutations) cells with a different phenotype randomly. Since then, there have been many further developments that incorporate stochastic elements into the model, such as those proposed by ***Lea and Coulson*** (***1949***), ***Koch*** (***1982***), ***Luebeck and Moolgavkar*** (***2002***), and ***Dewanji et al***. (***2005***). We can find various stochastic processes: Poisson processes (***Bartoszynski et al., 1981***), Markov chains (***Gupta et al., 2011***), and branching processes (***Jiang et al., 2017***), or even random sums of birth-death processes (***Dewanji et al., 2005***), all playing key roles in the mathematical theories of cellular clonal growth and evolution. These models have been applied to clinical data on lung cancer (***Newton et al., 2012***), breast cancer (***Speer et al., 1984***), and treatment of cancer (***Spina et al., 2014***).

At single-cell resolution, another cause for departure from exponential growth is the presence of positive (growth promoting) cell-cell interactions (Allee effect) in the early phase of population growth, such that cell density plays a role in stimulating division, giving rise to the critical mass dynamics (***Johnson et al., 2019; Korolev et al., 2014***).

To understand the intrinsic tumor growth behavior (change of tumor volume over time) it is therefore essential to study tumor cell populations in culture which affords detailed quantitative analysis of cell numbers over time, unaffected by the tumor microenvironment, and to measure departure from exponential growth. This paper focuses on stochastic growth of clonal but phenotypically heterogeneous HL60 leukemia cells with near single-cell sensitivities in the early phase of growth, that is, in sparse cultures. We and others have in the past years noted that at the level of single cells, each cell behaves akin to an individual, differently from another, which can be explained by the slow correlated transcriptome-wide fluctuations of gene expression (***Chang et al., 2008; Li et al., 2016***). Given the phenotypic heterogeneity and anticipated functional consequences, grouping of cells is necessary. Such classification would require molecular cell markers for said functional implication, but such markers are often di*Z*cult to determine a priori. Here, since most pertinent to cancer biology, we directly use a functional marker that is of central relevance for cancer: cell division, which maps into cell population growth potential *—* in brief *“*cell growth*”*.

Therefore, we monitored longitudinally the growth of cancer cell populations seeded at very small numbers of cells (1, 4, or 10 cells) in statistical ensembles of microcultures (wells on a plate of wells). We found evidence that clonal HL60 leukemia cell populations contain subpopulations that exhibit diverse growth patterns. Based on statistical analysis, we propose the existence of three distinctive cell phenotypic states with respect to cell growth. We show that a branching process model captures the population growth kinetics of a population with distinct cell subpopulations. Our results suggest that the initial phase cell growth (*“*take-off*”* of a cell culture) in the HL60 leukemic cells is predominantly driven by the fast-growing cell subpopulation. Reseeding experiments revealed that the fast-growing subpopulation could maintain its growth rate over several cell generations, even after the placement in a new environment. Our observations underscore the need to not only target the fast-growing cells but also the transition to them from the other cell subpopulations.

## Results

### Experiment of the cell population growth from distinct initial cell numbers

To expose the variability of growth kinetics as a function of initial cell density *N*_0_ (*“*initial seed number*”*), HL60 cells were sorted into wells of a 384-well plate (0.084 cm^2^ area) to obtain *“*statistical ensembles*”* of replicate microcultures (wells) of the same condition, distinct only by *N*_0_. Based on prior titration experiments to determine ranges of interest for *N*_0_ and statistical power, for this experiment we plated 80 wells with *N*_0_ = 10 cells (*N*_0_ = 10-cell group), 80 wells with *N*_0_ = 4 cells (*N*_0_ = 4-cell group), and 80 wells with *N*_0_ = 1 cell (*N*_0_ = 1-cell group). Cells were grown in the same conditions for 23 days (for details of cell culture and sorting, see the Methods section). Digital images were taken every 24 hours for each well from Day 4 on, and the area occupied by cells in each well was determined using computational image analysis. We had previously determined that one area unit equals approximately 500 cells. This is consistent and readily measurable because the relatively rigid and uniformly spherical HL60 cells grow as a non-adherent *“*packed*”* monolayer at the bottom of the well. Note that we are interested in the initial exponential growth (and departure from it) and not in the latter phases when the culture becomes saturated as has been the historical focus of analysis (see Introduction).

Wells that have reached at least 5 area units were considered for the characterization of early phase (before plateau) growth kinetics by plotting the areas in logarithmic scale as a function of time (Fig. 1). All the *N*_0_ = 10-cell wells required 3.6-4.6 days to grow from 5 area units to 50 area units (mean=4.05, standard deviation=0.23). For the *N*_0_ = 1-cell wells, we observed a diversity of behaviors. While some of the cultures only took 3.5-5 days to grow from 5 area units to 50 area units, others needed 6-7.2 days (mean=5.02, standard deviation=0.75). The *N*_0_ = 4-cell wells had a mean=4.50 days and standard deviation=0.44 to reach that same population size.

**Figure 1.**
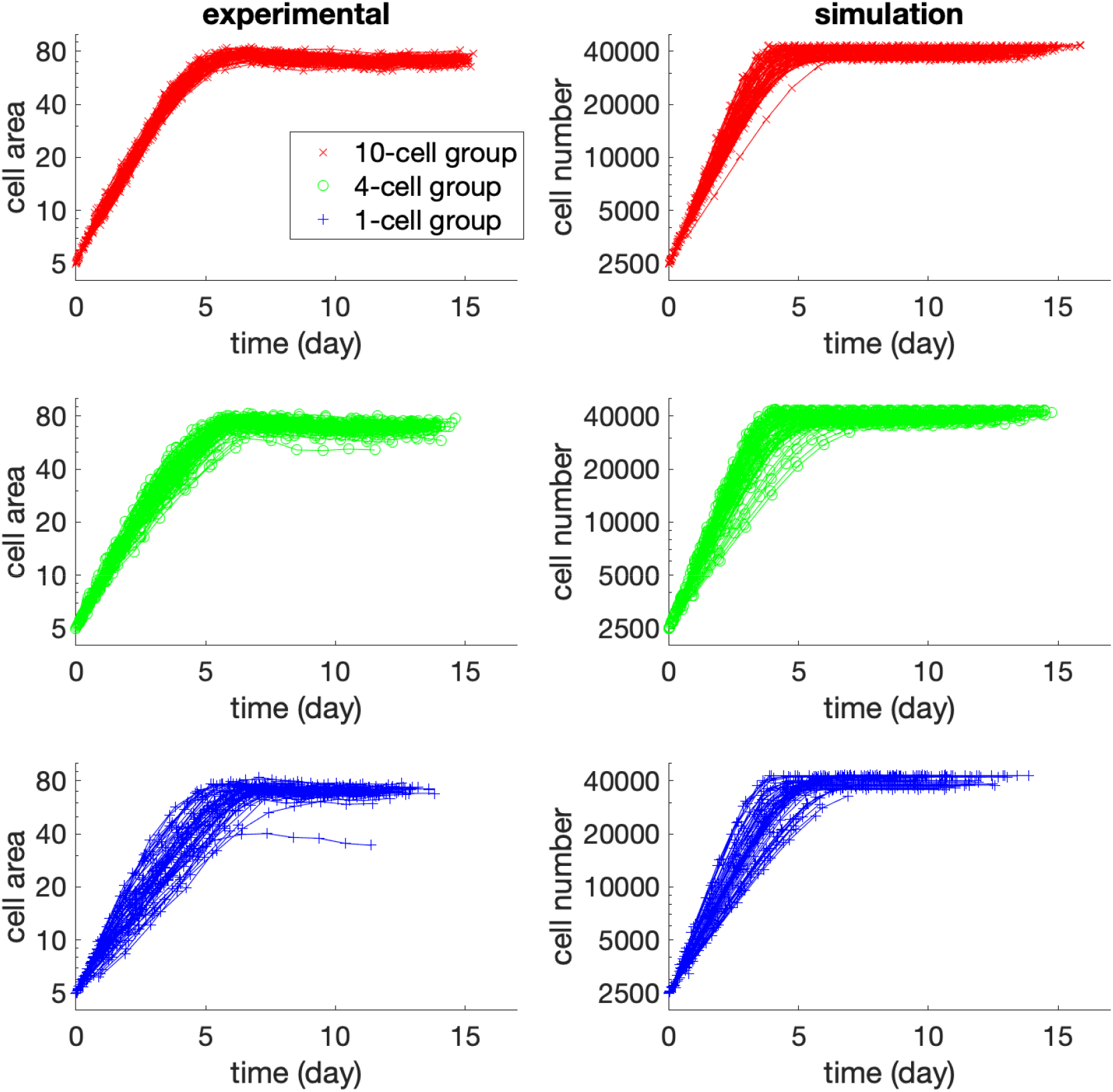
Growth curves of the experiment (left) and simulation (right), starting from the time of reaching 5 area units (experiment) or having 2500 cells (simulation), with a logarithm scale for the *y*-axis. The time required for reaching 5 area units was determined by exponential extrapolation, as reliable imaging started at *>* 5 area units. The *x*-axis is the time from reaching 5 area units (experiment) or 2500 cells (simulation). Red, green, or blue curves correspond to 10, 4, or 1 initial cell(s). Although starting from the same population level, patterns are different for distinct initial cell numbers. The *N*_0_ = 1-cell group has higher diversity.

To examine the exponential growth model, in Fig. 2 (left panel), we plotted the per capita growth rate versus cell population size, where each point represents a well (population) at a time point.

**Figure 2.**
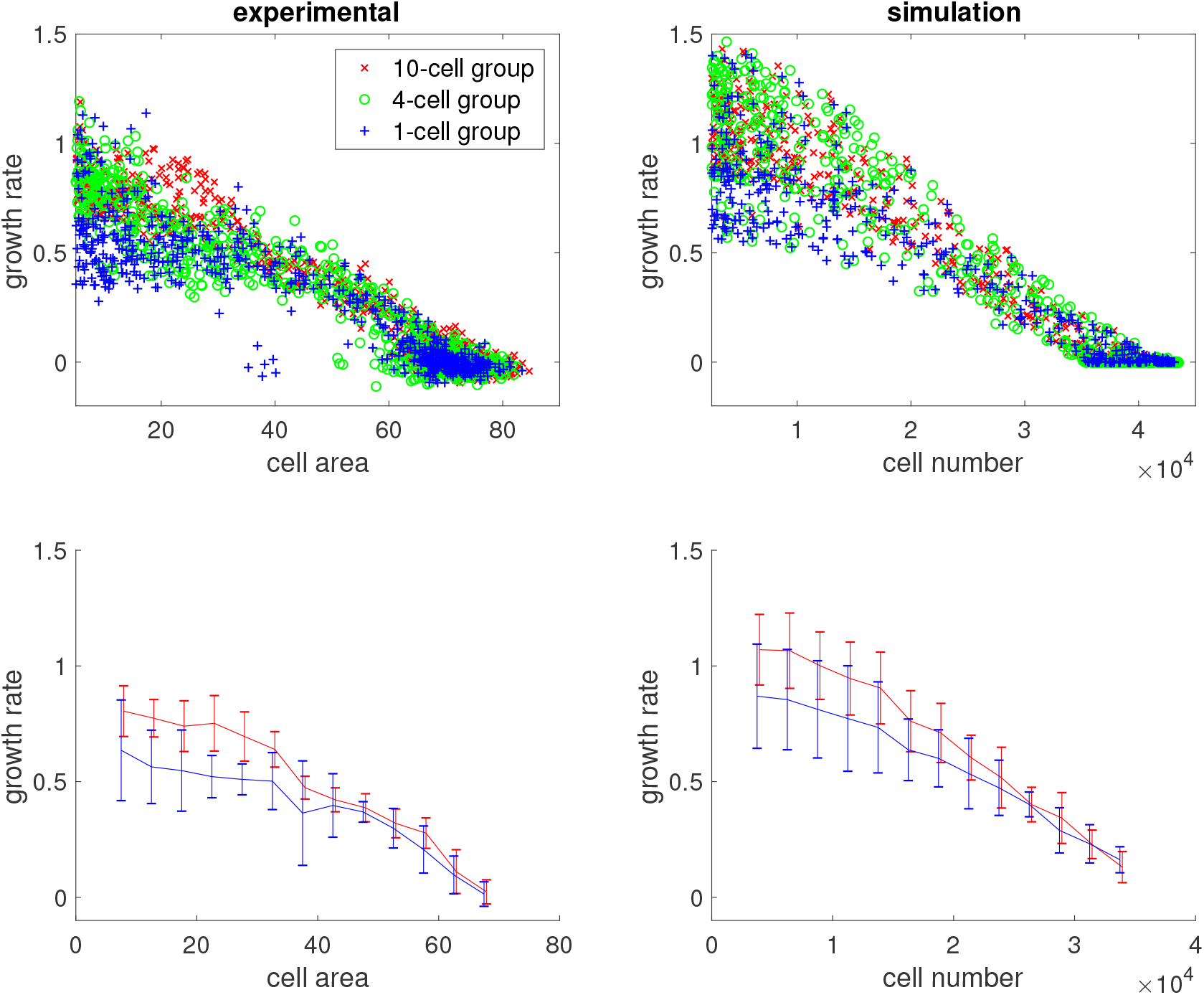
Per capita growth rate (averaged within one day) vs. cell population for the experiment (left) and simulation (right). Each point represents one well in one day. Red, green, or blue points correspond to 10, 4, or 1 initial cell(s).

As expected, as the population became crowded, the growth rate decreased toward zero. But in the earlier phase, many populations in the *N*_0_ = 1-cell group had a lower per capita growth rate than those in the *N*_0_ = 10-cell group, even at the same population size – thus departing from the expected behavior of exponential growth. The weighted Welch*’*s *t*-test showed that the difference in these growth rates was significant (see the Methods section).

While qualitative differences in the behaviors of cultures with different initial seeding cell numbers *N*_0_ can be expected for biological reasons (see below), in the elementary exponential growth model, the difference of growth rate should disappear when populations with distinct seeding numbers are aligned for the same population size that they have reached as in Fig. 2. A simple possibility is that the deviations of expected growth rates emanate from difference in cellintrinsic properties. Some cells grew faster, with a per capita growth rate of 0.6 ∼ 0.9 (all *N*_0_ = 10-cell wells and some *N*_0_ = 1-cell wells), while some cells grew slower, with a per capita growth rate of 0.3 ∼ 0.5 (some of the *N*_0_ = 1-cell wells). In other words, there is intrinsic heterogeneity in the cell population that is not *“*averaged out*”* in the culture with low *N*_0_, and the sampling process exposes these differences between the cells that appear to be relatively stable.

To illustrate the inherent diversity of initial growth rates, in Fig. 3 (left panel), we display the daily cell-occupied areas plotted on a linear scale starting from Day 4. All wells with seed of *N*_0_ = 10 or *N*_0_ = 4 cells grew exponentially. Among the *N*_0_ = 1-cell wells, 14 populations died out. Four wells in the *N*_0_ = 1-cell group had more than 10 cells on Day 8 but never grew exponentially, and had fewer than 1000 cells after 15 days (on Day 23). For these non-growing or slow-growing *N*_0_ = 1-cell wells, the per capita growth rate was 0 ∼ 0.2. In comparison, all the *N*_0_ = 10-cell wells needed at most 15 days to reach the carrying capacity (around 80 area units, or 40000 cells). See Table 1 for a summary of the *N*_0_ = 1-cell group*’*s growth patterns. This behavior is not idiosyncratic to the culture system because they recapitulate a pilot experiment performed in the larger scale format of 96-well plates (not shown).

**Table 1.**
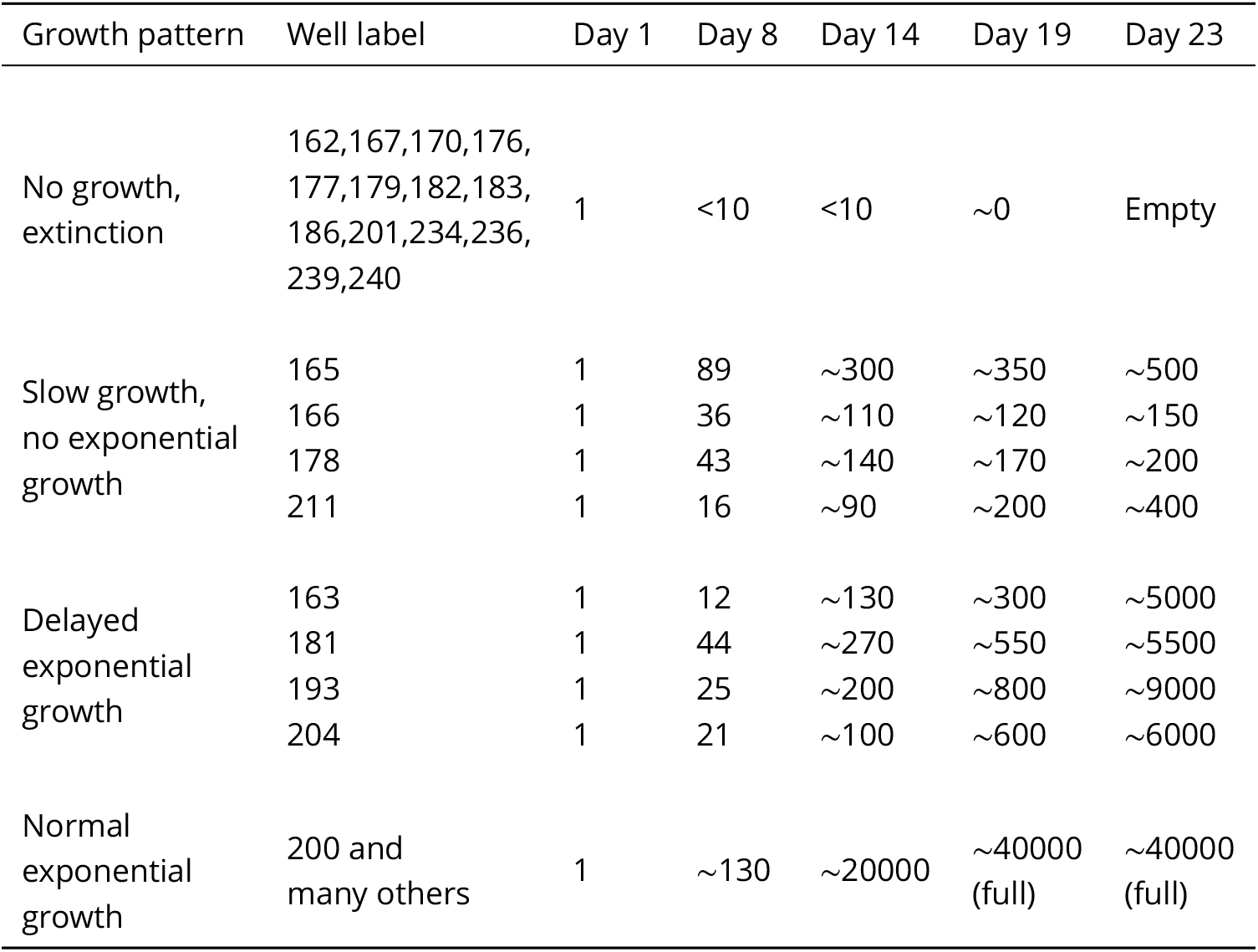
The population of some wells in the *N*_0_ = 1-cell group in the growth experiment with different initial cell numbers, where ∼ meant approximate cell number. These wells illustrated different growth patterns from those wells starting with *N*_0_ = 10 or *N*_0_ = 4 cells. Such differences implied that cells from wells with different initial cell numbers were essentially different.

| Growth pattern | Well label | Day 1 | Day 8 | Day 14 | Day 19 | Day 23 |
| --- | --- | --- | --- | --- | --- | --- |
| No growth, extinction | 162,167,170,176, 177,179,182,183, 186,201,234,236, 239,240 | 1 | <10 | <10 | ~0 | Empty |
| Slow growth, no exponential growth | 165 | 1 | 89 | ~300 | ~350 | ~500 |
|  | 166 | 1 | 36 | ~110 | ~120 | ~150 |
|  | 178 | 1 | 43 | ~140 | ~170 | ~200 |
|  | 211 | 1 | 16 | ~90 | ~200 | ~400 |
| Delayed exponential growth | 163 | 1 | 12 | ~130 | ~300 | ~5000 |
|  | 181 | 1 | 44 | ~270 | ~550 | ~5500 |
|  | 193 | 1 | 25 | ~200 | ~800 | ~9000 |
|  | 204 | 1 | 21 | ~100 | ~600 | ~6000 |
| Normal exponential growth | 200 and many others | 1 | ~130 | ~20000 | ~40000 (full) | ~40000 (full) |

**Figure 3.**
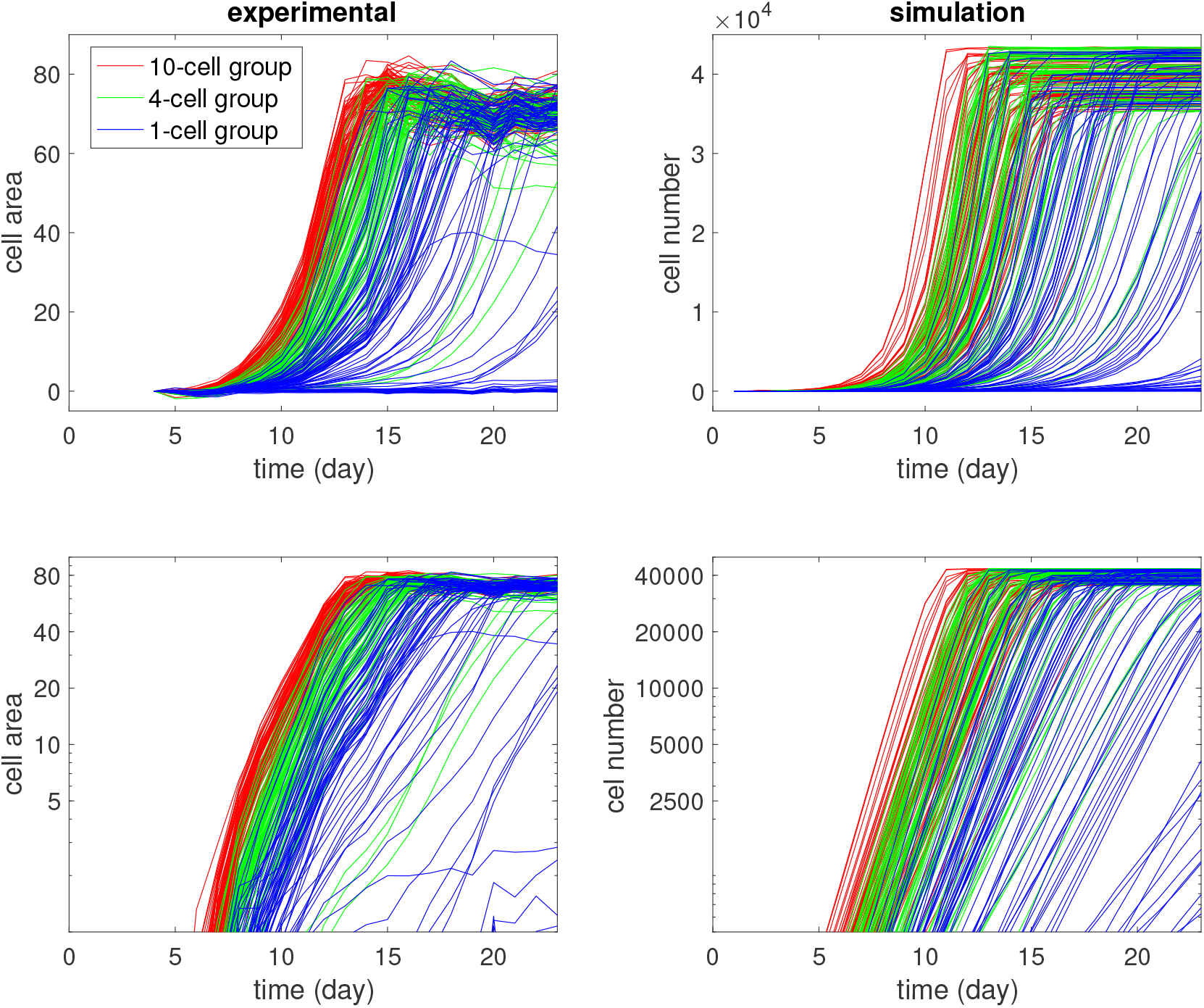
Growth curves of the experiments with different initial cell numbers *N*_0_ (left) and growth curves of corresponding simulation (right). Each curve describes the change in the cell population (measured by area or number) over a well along time. Red, green, or blue curves correspond to *N*_0_ = 10, 4, or 1 initial cell(s).

From the above experimental observations, we asserted that there might be at least three stable cell growth phenotypes in a population: a fast type, whose growth rate was 0.6 ∼ 0.9/day for non-crowded conditions; a moderate type, whose growth rate was 0.3 ∼ 0.5/day for non-crowded conditions; and a slow type, whose growth rate was 0 ∼ 0.2/day for the non-crowded population.

The graphs of Fig. 3 also revealed other phenomena of growth kinetics: (1) Most *N*_0_ = 4-cell wells plateaued by Day 14 to Day 17, but some lagged significantly behind. (2) Similarly, four wells in the *N*_0_ = 1-cell group exhibited longer lag-times before the exponential growth phase, and never reached half-maximal cell numbers by Day 23. These outliers reveal intrinsic variability and were taken into account in the parameter scanning (see the Methods section).

### Reseeding experiments revealing the enduring intrinsic growth patterns

When a well in the *N*_0_ = 1-cell group had grown to 10 cells, population behavior was still different from those in the *N*_0_ = 10-cell group at the outset. In view of the spate of recent results revealing phenotypic heterogeneity, we hypothesized that the difference was cell-intrinsic as opposed to being a consequence of the environment (e.g., culture medium in *N*_0_ = 1 vs *N*_0_ = 10 -cell wells).

To test our hypothesis and exclude differences in the culture environment as determinants of growth behavior, we reseeded the cells that exhibited the different growth rates in fresh cultures. We started with a number of *N*_0_ = 1-cell wells. After a period of almost 3 weeks, again some wells showed rapid proliferation, with cells covering the well, while others were half full and yet others wells were almost empty. We collected cells from the full and half-full wells and reseeded them into 32 wells each (at about *N*_0_ = 78 cells per well). These 64 wells were monitored for another 20 days. We found that most wells reseeded from the full well took around 11 days to reach the population size of a half-full well, while most wells reseeded from the half-full well required around 16 ∼ 20 days to reach the same half full well population size. Five wells reseeded from the half-full wells were far from even reaching half full well population size by Day 20 (see Table 2). Permutation test showed that this difference in growth rate was significant (see the Methods section).

**Table 2.**
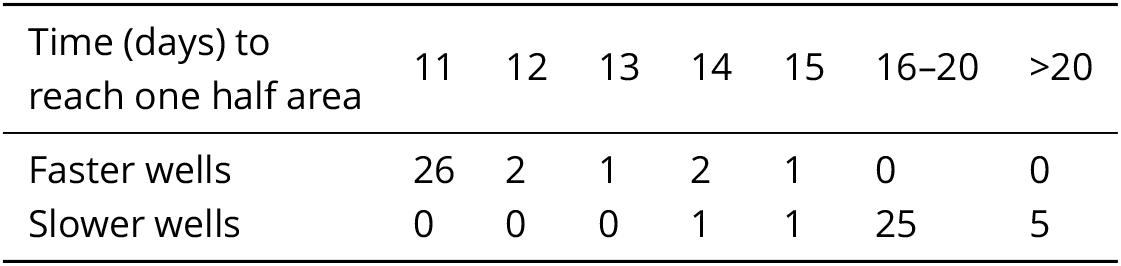
The distribution of time needed for each well to reach the *“*half area*”* population size in the reseeding experiment. We reseeded equal numbers of cells that grew faster (from a full well) and cells that grew slower (from a half-full well), and cultivated them under the same new fresh medium environment to compare their intrinsic growth rates. The results showed that faster growing cells, even reseeded, still grew faster.

This reseeding experiment shows that the difference in growth rate was maintained over multiple generations, even after slowing down in the plateau phase (full well) and was maintained when restarting a microculture at low density in fresh medium devoid of secreted cell products. Therefore, it is plausible that there exists endogenous heterogeneity of growth phenotypes in the clonal HL60 cell line and that these distinct growth phenotypes are stable for at least 15 ∼ 20 cell generations.

### Quantitative analysis of experimental results

In the experiments with different initial cell numbers *N*_0_, we observed at least three patterns with different growth rates, and the reseeding showed that these growth patterns were endogenous to the cells. Therefore, we propose that each growth pattern discussed above corresponded to a cell phenotype that dominated the population: fast, moderate, and slow.

In the initial seeding of cells that varies *N*_0_, the cells were randomly chosen (by FACS); thus, their intrinsic growth phenotypes were randomly distributed. During growth, the population of a well would be dominated by the fastest type that existed in the seeding cells, thus qualitatively, we have following scenarios: (1) A well in the *N*_0_ = 10-cell group almost certainly had at least one initial cell of fast type, and the population would be dominated by fast type cells. Different wells had almost the same growth rate, reaching saturation at almost the same time. (2) For an *N*_0_ = 1-cell well, if the only initial cell is of the fast type, then the population has only the fast type, and the growth pattern will be close to that of *N*_0_ = 10-cell wells. If the only initial cell is of the moderate type, then the population could still grow exponentially, but with a slower growth rate. This explains why after reaching 5 area units, many but not all *N*_0_ = 1-cell wells were slower than *N*_0_ = 10-cell wells. (3) Moreover, in such an *N*_0_ = 1-cell well with a moderate type initial cell, the cell might not divide quite often during the first few days due to randomness of entering the cell cycle. This would lead to a considerable delay in entering the exponential growth phase. (4) By contrast, for an *N*_0_ = 1-cell well with a slow type initial cell, the growth rate could be too small, and the population might die out or survive without ever entering the exponential growth phase in duration of the experiment. (5) Most *N*_0_ = 4-cell wells had at least one fast type initial cell, and the growth pattern was the same as *N*_0_ = 10-cell wells. A few *N*_0_ = 4-cell wells only had moderate and slow cells, and thus had slower growth patterns.

The above verbal argument is shown in Fig. 4 and entails mathematical modeling with the appropriate parameters that relate the relative frequency of these cell types in the original population, their associated growth and transition rates to examine whether it explains the data.

**Figure 4.**
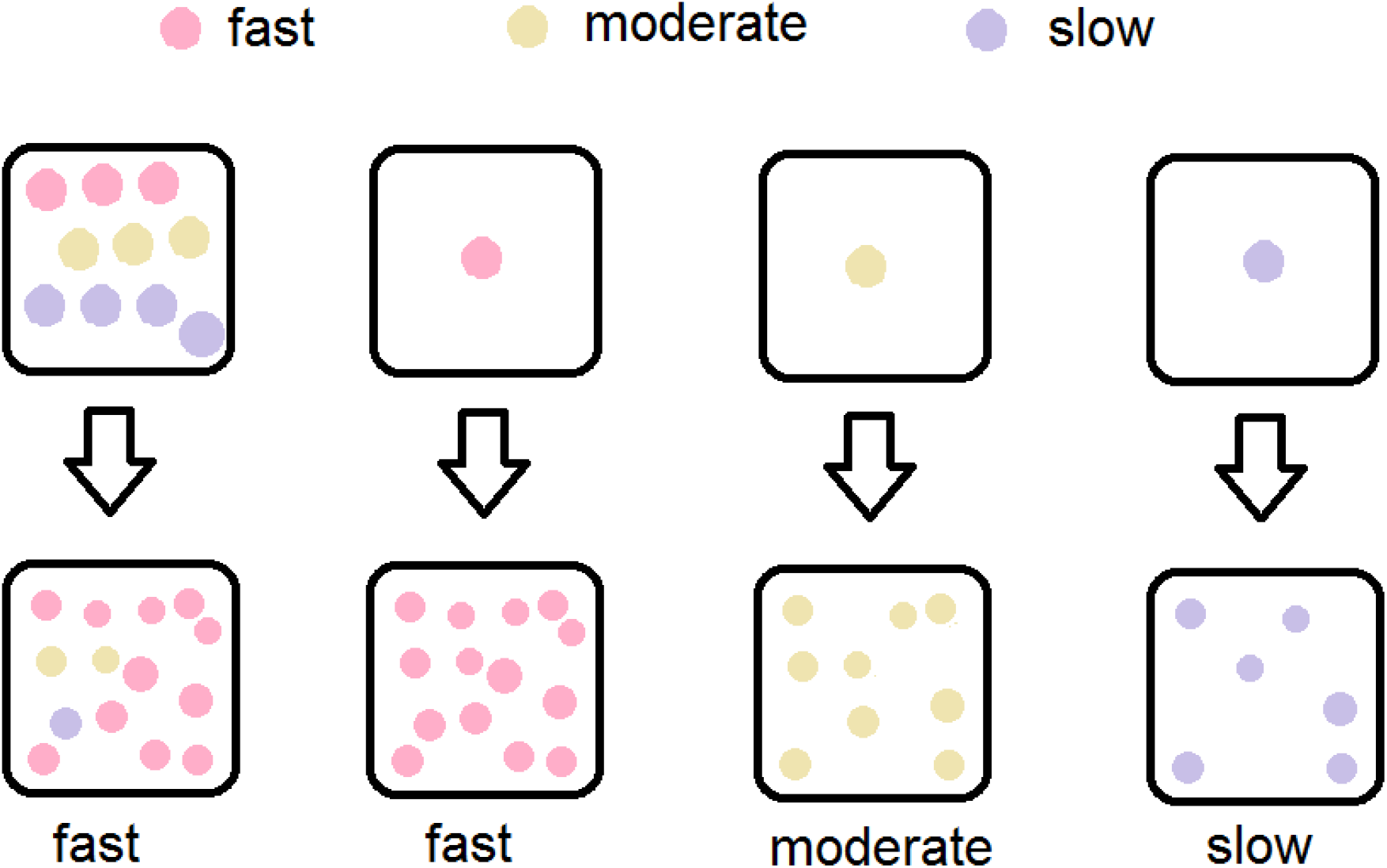
Schematic illustration of the qualitative argument: Three cell types and growth patterns (three colors) with different seeding numbers. One *N*_0_ = 10-cell well will have at least one fast type cell with high probability, which will dominate the population. One *N*_0_ = 1-cell well can only have one cell type, thus in the microculture ensemble of replicate wells, three possible growth patterns for wells can be observed.

### Branching process model

To construct a quantitative dynamical model to recapitulate the growth dynamics differences from cell populations with distinct initial seed cell numbers *N*_0_, and three intrinsic types of proliferation behaviors, we used a multi-type discrete-time branching process.

The traditional method of population dynamics based on ordinary differential equation (ODE), which is deterministic and has continuous variables, is not suited when the cell population is small as is the case for the earliest stage of proliferation from a few cells being studied in our experiments. Deterministic models are also unfit because with such small populations and measurements at single-cell resolution, stochasticity in cell activity does not average out. The nuanced differences between individual cells cannot be captured by a different deterministic mechanism of each individual cell, and the only information available is the initial cell number. Thus, the unobservable nuances between cells are taken care of by a stochastic model.

Given the small populations, our model should be purely stochastic, without deterministic growth. The focus is the concrete population size of a finite number (three) of types, thus Poisson processes are not suitable. Markov chains can partially describe the proportions under some conditions, but population sizes are known, not just their ratios, therefore Markov chains are not necessary. Branching processes can describe the population size of multiple types with symmetric and asymmetric division, transition, and death (***Jiang et al., 2017***). Also, the parameters can be temporally and spatially inhomogeneous, which is convenient. Therefore, we utilized branching processes in our model.

In the branching process, each cell during each time interval independently and randomly chooses a behavior: division, death, or stagnation in the quiescent state, whose rates depend on the cell growth type. Denoting the growth rate and death rate of the fast type by *g*_F_ and _F_ respectively, and the population size of fast type cells on Day *n* by *F* (*n*), the population at Day *n* + 1 is:

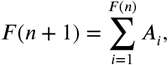

where *A*_*i*_ for different *i* are independent. *A*_*i*_ represents the descendants of a fast type cell *i* after one day. It equals 2 with probability *g*_F_, 0 with probability _F_, and 1 with probability 1 − *g*_F_ − _F_. Therefore, given *F* (*n*), the distribution of *F* (*n* + 1) is:

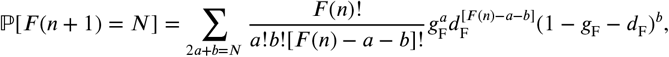

where the summation is taken for all non-negative integer pairs (*a, b*) with 2*a* + *b* = *N*. Moderate and slow types evolve similarly, with their corresponding growth rates *g*_M_, *g*_S_, and death rates _M, S_.

As shown in Fig. 2, the growth rates *g*_F_, *g*_M_, and *g*_S_ should be decreasing functions of the total population. In our model, we adopted a quadratic function.

We performed a parameter scan to show that our model could reproduce experimental phenomena for a wide range of model parameters (see details in Table 3).

**Table 3.**
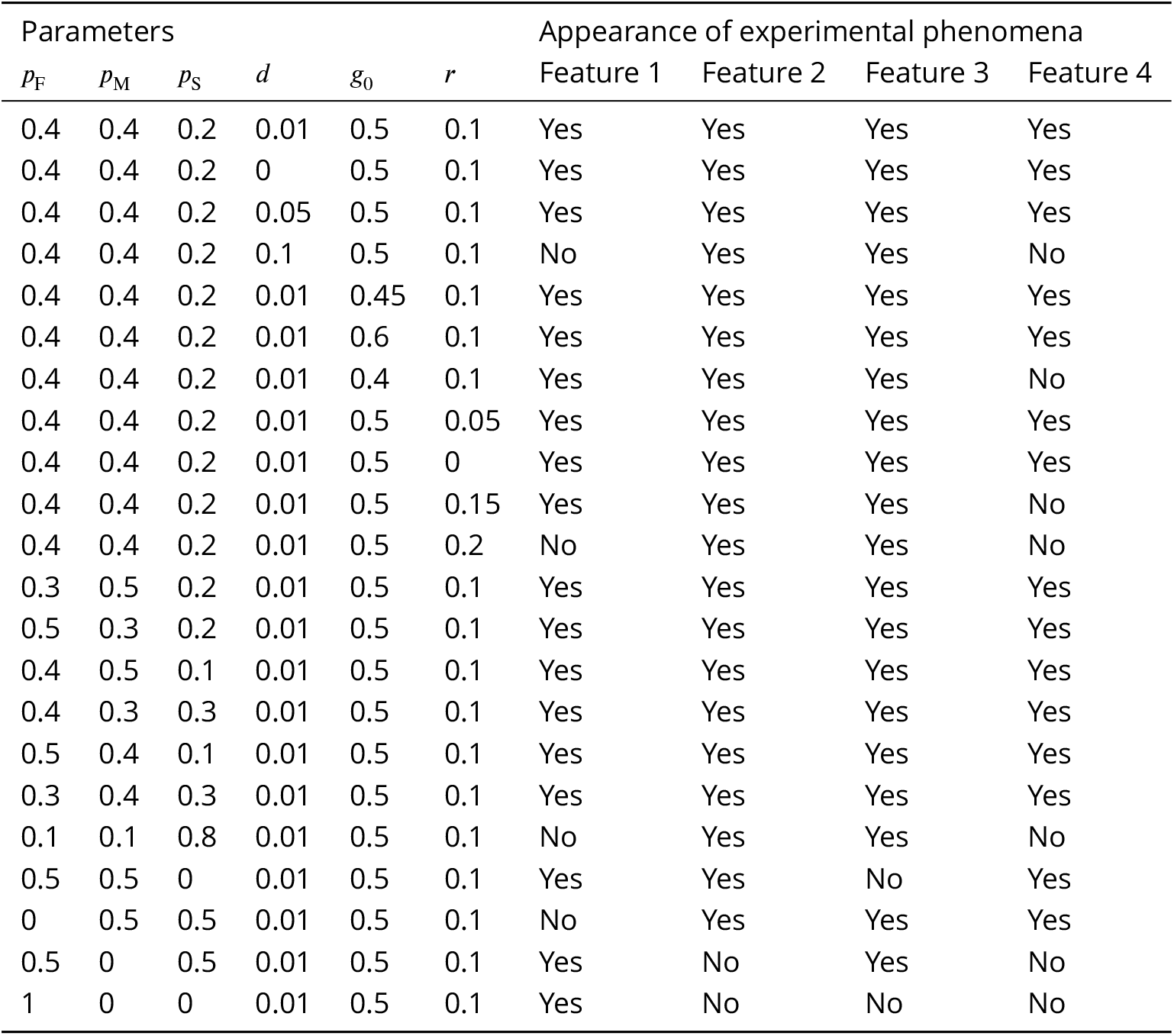
Performance of our model with different parameters. Here we adjusted the parameters of our model in a wide range and observed whether the model could still reproduce four important *“*features*”* in the experiment. This parameter scan showed that our model is robust under perturbations on parameters. Here *p*_F_, *p*_M_, *p*_S_ are the probabilities that an initial cell is of fast, moderate, or slow type; is the death rate; *g*_0_ is the growth factor; *r* is the range of the random modi*1*er. See the Methods section for explanations of these parameters. Feature 1, all wells in the *N*_0_ = 10-cell group were saturated; Feature 2, presence of late-growing wells in the *N*_0_ = 1-cell group; Feature 3, presence of non-growing wells in the *N*_0_ = 1-cell group; Feature 4, different growth rates at the same population size between the *N*_0_ = 10-cell group and the *N*_0_ = 1-cell group.

The simulation results are shown on the right panels of Figs. 1–3, in comparison with the experimental data in the left. Our model qualitatively captured the growth patterns of groups with different initial seeding cell numbers. For example, in Fig. 2, when wells were less than half full (cell number *<* 20000), most wells in the *N*_0_ = 10-cell group grew faster than the *N*_0_ = 1-cell group even when they had the same cell number. In Fig. 3, all wells in the *N*_0_ = 10-cell group in our model grew quickly until saturation. Similar to the experiment, some wells in the *N*_0_ = 1-cell group in our model never grew, while some began to take off very late.

In our model, the high extinction rate in the *N*_0_ = 1-cell group (14/80) was explained as *“*bad luck*”* at the early stage, since birth rate and death rate were close, and a cell could easily die without division. Another possible explanation for such a difference in growth rates was that the population would be 10 small colonies when starting from 10 initial cells, while starting from 1 initial cell, the population would be 1 large colony. With the same area, 10 small colonies should have a larger total perimeter, thus larger growth space and larger growth rate than that of 1 large colony. However, we carefully checked the photos, and found that almost all wells produced 1 large colony with nearly the same shape, and there was no significant relationship between colony perimeter and growth rate.

## Discussion

As many recent single-cell level data have shown, a tumor can contain multiple distinct subpopulations engaging in interconversions and interactions among them that can influence cancer cell proliferation, death, migration, and other features that contribute to malignancy (***Pisco and Huang, 2015; Zhou et al., 2014a; Angelini et al., 2022; Howard et al., 2018; Sahoo et al., 2021; Zhou et al., 2014b; Johnson et al., 2019; Korolev et al., 2014; Chapman et al., 2014; Niu et al., 2015; Chen et al., 2016***). Presence of these two intra-population behaviors can be manifest as departure from the elementary model of exponential growth (***Skehan and Friedman, 1984***) (in the early phase of population growth, far away from carrying capacity of the culture environment which is trivially non-exponential). The exponential growth model assumes uniformity of cell division rates across all cells (hence a population doubling rate that is proportional to a given population size *N*(*t*)) and the absence of cell-cell interactions that affect cell division and death rates. Investigating the *“*non-genetic heterogeneity*”* hypothesis of cancer cells quantitatively is therefore paramount for understanding cancer biology but also for elementary principles of cell population growth.

As an example, here we showed that clonal cell populations of the leukemia HL60 cell line are heterogeneous with regard to growth behaviors of individual cells that can be summarized in subpopulations characterized by a distinct intrinsic growth rates which were revealed by analysis of the early population growth starting with microcultures seeded with varying (low) cell number *N*_0_. Since we have noted only very weak effect of cell-cell interactions on cell growth behaviors (Allee effect) in this cell line (as opposed to another cell tumor cell line in which we found that departure from exponential growth could be explained by the Allee effect (***Johnson et al., 2019***)), we focused on the very presence among HL60 cells of subpopulations with distinct proliferative capacity as a mechanism for the departure of the early population growth curve from exponential growth.

The reseeding experiment demonstrated that the characteristic growth behaviors of subpopulations could be inherited across cell generations and after moving to a new environment (fresh culture), consistent with long-enduring endogenous properties of the cells. This result might be explained by cells occupying distinct stable cell states (in a multi-stable system). Thus, we introduced multiple cell types with different growth rates in our stochastic model. Specifically, in a branching process model, we assumed the existence of three types: fast, moderate, and slow cells. The model we built could replicate the key features in the experimental data, such as different growth rates at the same population size between the *N*_0_ = 10-cell group and the *N*_0_ = 1-cell group, and the presence of late-growing and non-growing wells in the *N*_0_ = 1-cell group.

While we were able to fit the observed behaviors in which the growth rate depended not only on *N*(*t*) but also on *N*_0_, the existence of the three or even more cell types still needs to be verified experimentally. For instance, statistical cluster analysis of transcriptomes of individual cells by single-cell RNA-seq (***Bhartiya et al., 2021***) over the population may identify the presence of transcriptomically distinct subpopulations that could be isolated (e.g., after association with cell surface markers) and evaluated separately for their growth behaviors.

The central assumption of coexistence of multiple subpopulations in the cell line stock must be accompanied by the second assumption that there are transitions between these distinct cell populations. For otherwise, in the stock population the fastest growing cell would eventually outgrow the slow growing cells. Furthermore, one has to assume a steady-state in which the population of slow growing cells are continuously replenished from the population of fast-growing cells. Finally, we must assume that the steady-state proportions of the subpopulations are such that at low seeding wells with *N*_0_ = 1 cells, there is a sizable probability that a microculture receives cells from each of the (three) presumed subtypes of cells. The number of wells in the ensemble of replicate microcultures for each *N*_0_condition has been su*Z*ciently large for us to make the observations and inform the model, but a larger ensemble would be required to determine with satisfactory accuracy the relative proportions of the cell types in the parental stock population.

Transitions might also have been happening during our experiment. For example, those late growing wells in the *N*_0_ = 1-cell group could be explained by such a transition: Initially, only slow type cells were present, but once one of these slow growing cells switched to the moderate type, an exponential growth ensued at the same rate that is intrinsic to that of moderate cells.

If there are transitions, what is the transition rate? Our reseeding experiments are compatible with a relatively slow rate for interconversion of growth behaviors in that the same growth type was maintained across 30 generations. An alternative to the principle of transition at a constant intrinsic to each of the types of cells may be that transition is extrinsically determined. Specifically, the seeding in the *“*lone*”* condition of *N*_0_ = 1 may *induce* a dormant state, that is a transition to a slower growth mode that is then maintained, on average over 30+ generations, with occasional return to the faster types that account for the delayed exponential growth.

This model however would bring back the notion of *“*environment awareness*”*, or the principle of a *“*critical density*”* for growth implemented by cell-cell interaction (Allee effect) which we had deliberately not considered (see above) since it was not necessary. We do not exclude this possibility which could be experimentally tested as follows: Cultivate *N*_0_ = 1-cell wells for 20 days when the delayed exponential growth has happened in some wells, but then use the cells of those wells with fast-growing population (which should contain of the fast type) to restart the experiment, seeded at *N*_0_ = 10, 4, 1 cells. If wells with different seeding numbers exhibit the same growth rates, then the growth difference in the original experiment is solely due to preexisting (slow interconverting) cell phenotypes. If now the *N*_0_ = 1-cell wells resumes the typical slow growth, this would indicate a density induced transition to the slow growth type.

In the spirit of Occam*’*s razor, and given the technical di*Z*culty in separate experiments to demonstrate cell-cell interactions in HL60 cells, we were able to model the observed behaviors with the simplest assumption of cell-autonomous properties, including existence of multiple states (growth behaviors) and slow transitions between them but without cell density dependence or interactions.

Taken together, we showed that one manifestation of the burgeoning awareness of ubiquitous cell phenotype heterogeneity in an isogenic cell population is the presence of distinct intrinsic types of cells that slowly interconvert among them, resulting in a stationary population composition. The differing growth rates of the subtypes and their stable proportions may be an elementary characteristic of a given population that by itself can account for the departure of early population growth kinetics from the basic exponential growth model.

## Methods

### Setup of growth experiment with different initial cell numbers

HL60 cells were maintained in IMDM wGln, 20% FBS(heat inactivated), 1% P/S at a cell density between 3 × 10^5^ and 2.5 × 10^6^ cells/ml (GIBCO). Cells were always handled and maintained under sterile conditions (tissue culture hood; 37°C, 5% CO_2_, humidified incubator). At the beginning of the experiment, cells were collected, washed two times in PBS, and stained for vitality (Trypan blue GIBCO). The population of cells was first gated for morphology and then for vitality staining. Only Trypan negative cells were sorted (BD FACSAria II). The cells were sorted in a 384 well plate with IMDM wGln, 20% FBS(heat inactivated), and 1% P/S (GIBCO).

Cell population growth was monitored using a Leica microscope (heated environmental chamber and CO_2_ levels control) with a motorized tray. Starting from Day 4, the 384 well plate was placed inside the environmental chamber every 24 hours. The images were acquired in a 3 × 3 grid for each well; after acquisition, the 9 fields were stitched into a single image. Software ImageJ was applied to identify and estimate the area occupied by *“*entities*”* in each image. The area (proportional to cell number) was used to follow the cell growth.

### Setup of reseeding experiment for growth pattern inheritance

HL60 cells were cultivated for 3 weeks, and then we chose one full well and one half full well. We supposed the full well was dominated by fast type cells, and the half-full well was dominated by moderate type cells, which had lower growth rates. We reseeded cells from these two wells and cultivated them in two 96-well (rows A-H, columns 1-12) plates. In each plate, B2-B11, D2-D11, and F2-F11 wells started with 78 fast cells, while C2-C11, E2-E11, and G2-G11 wells started with 78 moderate cells. Rows A, H, columns 1, 12 had no cells and no media, and we found that wells in rows B, G, columns 2, 11, which were the outmost non-empty wells, evaporated much faster than inner wells. Therefore, the growth of cells in those wells was much slower than inner wells. Hence we only considered inner wells, where D3-D10 and F3-F10 started with fast cells, C3-C10 and E3-E10 started with moderate cells, namely 32 fast wells and 32 moderate wells in total. During the experiment, no media was added. Each day, we observed those wells to check whether their areas exceeded one-half of the whole well. The experiment was terminated after 20 days.

### Weighted Welch*’*s *t*-test

The weighted Welch*’*s *t*-test is used to test the hypothesis that two populations have equal mean, while sample values have different weights (***Goldberg et al., 2005***). Assume for group *i* (*i* = 1, 2), the sample size is *N*_*i*_ and the *j*th sample is the average of 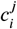 independent and identically distributed variables. Let 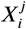 be the observed average for the *j*th sample. Set *v*_1_ = *N*_1_ − 1, *v*_2_ = *N*_2_ − 1. Define

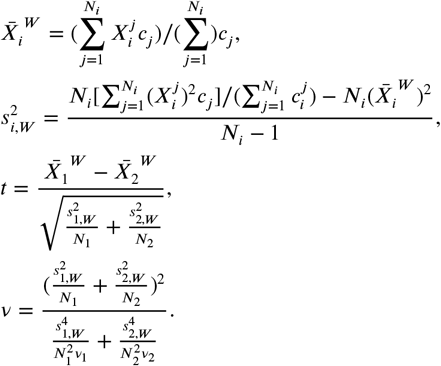

If two populations have equal mean, then *t* satisfies the *t*-distribution with degree of freedom *v*.

The weighted Welch*’*s *t*-test was applied to the growth experiment with different initial cell numbers, in order to determine whether the growth rates during exponential phase (5–50 area units) were different between groups. Here 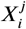 corresponded to growth rate, and 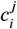 corresponded to cell area. The *p*-value for *N*_0_ = 10-cell group vs. *N*_0_ = 4-cell group was 2.12 × 10^−8^; the *p*-value for *N*_0_ = 10-cell group vs. *N*_0_ = 1-cell group was smaller than 10^−12^; the *p*-value for *N*_0_ = 4-cell group vs. *N*_0_ = 1-cell group was 5.35 × 10^−5^. Therefore, the growth rate difference between any two groups was statistically significant.

### Permutation Test

The permutation test is a non-parametric method to test whether two samples are significantly different with respect to a statistic (e.g., sample mean) (***Hastie et al., 2016***). It is easy to calculate and fits our situation, thus we adopt this test rather than other more complicated tests, such as the Mann-Whitney test. For two samples {*x*_1_,., *x*_*m*_}, {*y*_1_,., *y*_*n*_}, consider the null hypothesis: the mean of *x* and *y* are the same. For these samples, calculate the mean of the first sample: 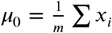. Then we randomly divide these *m* + *n* samples into two groups with size *m* and 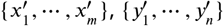, such that each permutation has equal probability. For these new samples, calculate the mean of the first sample: 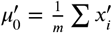. Then the two-sided *p*-value is defined as

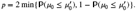

If *μ*_0_ is an extreme value in the distribution of 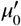, then the two sample means are different.

In the reseeding experiment, the mean time of exceeding half well for the fast group was 11.4375 days. For all 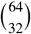 possible result combinations, only 7 combinations had equal or less mean time. Thus the *p*-value was 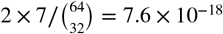. This indicated that the growth rate difference between fast group and moderate group was significant.

### Model Details

The simulation time interval was half day, but we only utilized the results in full days. For each initial cell, the probabilities of being fast, moderate or slow type, *p*_F_, *p*_M_, *p*_S_, were 0.4, 0.4, 0.2.

Each half day, a fast type cell had probability to die, and probability *g*_F_ to divide. The division produced two fast cells, capturing the intrinsic growth behavior that is to some extent inheritable. Denote the total cell number of previous day as *N*, then

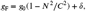

where *δ* is a random variable that satisfies the uniform distribution on [−*r, r*], and it is a constant for all cells in the same well. If *g*_F_ *<* 0, set *g*_F_ = 0. If *g*_F_ *>* 1 −, set *g*_F_ = 1 − *d*.

In the simulation displayed, death rate = 0.01, carrying capacity *C* = 40000, growth factor *g*_0_ = 0.5, and the range of random modifier *r* = 0.1.

Each half day, a moderate type cell had probability to die, and probability *g*_M_ to divide. The division produced two moderate cells. *g*_M_ = *g*_F_/1.5.

Similarly, each half day, a slow type cell had probability to die, and probability *g*_S_ to divide. The division produced two slow-growing cells. *g*_S_ = *g*_F_/3.

### Parameter scan

Since growth is measured by the area covered by cells, we could not experimentally verify most assumptions of our model, or determine the values of parameters. Therefore, we performed a parameter scan by evaluating the performance of our model for different sets of parameters. We adjusted 6 parameters: initial type probabilities *p*_F_, *p*_M_, *p*_S_, death rate, growth factor *g*_0_, and random modifier *r*. We checked whether these 4 features observable in the experiment could be reproduced: growth of all wells in the *N*_0_ = 10-cell group to saturation; existence of late-growing wells in the *N*_0_ = 1-cell group; existence of non-growing wells in the *N*_0_ = 1-cell group; difference in growth rates in the *N*_0_ = 10-cell group and the *N*_0_ = 1-cell group at the same population size. Table 3 shows the results of the performance of simulations with the various parameter sets. Within a wide range of parameters, our model is able to replicate the experimental results shown in Figs. 1–3, indicating that our model is robust under perturbations.

## Acknowledgments

This research was partially supported by NIGMS NIH-R01CA226258-01. We would like to thank Ivana Bozic, Yifei Liu, Georg Luebeck, Weili Wang, Yuting Wei and Lingxue Zhu for helpful advice and discussions.

## Data availability

The experimental data, simulation data, and corresponding code files could be found at https://github.com/YueWangMathbio/Leukemia.

